# Evolution of kinase polypharmacology across HSP90 drug discovery

**DOI:** 10.1101/2020.09.09.288936

**Authors:** Albert A. Antolin, Paul A. Clarke, Ian Collins, Paul Workman, Bissan Al-Lazikani

## Abstract

Most small molecules interact with several target proteins but this polypharmacology is seldom comprehensively investigated or explicitly exploited during drug discovery. Here, we use computational and experimental methods to systematically characterize the kinase cross-pharmacology of representative HSP90 inhibitors. We demonstrate that the resorcinol clinical candidates ganetespib and, to a lesser extent, luminespib, display unique off-target kinase pharmacology as compared to other HSP90 inhibitors. We also demonstrate that polypharmacology evolved during the optimisation to discover luminespib and that the hit, leads and clinical candidate all have different polypharmacological profiles. We conclude that the submicromolar target inhibition of protein kinases by ganetespib may have potential clinical significance and we recommend the computational and experimental characterization of polypharmacology earlier in drug discovery projects to unlock new multi-target drug design opportunities.

## Introduction

It is widely accepted that most small molecules will interact with multiple molecular targets when exposed to complex biological systems and the term polypharmacology is commonly used to refer to this phenomenon.^1,2^ It is also commonly acknowledged that off-targets can influence both drug efficacy and safety in the clinic.^3,4^ Moreover, polypharmacology of hit and lead compounds may inadvertently influence the direction of drug discovery projects. Despite this, polypharmacology is not being routinely explored as part of the drug discovery journey. Rather, to maximize cost-efficiency and maintain focus, potential off-targets are tested only for a very reduced number of compounds at late project stages.

Heat shock proteins (HSPs) are a group of molecular chaperones that are upregulated under stress to enable the correct folding of proteins.^5,6^ The 90 kDa heat shock protein HSP90 is one of the most abundant HSPs and a key regulator of proteostasis in both physiological conditions and under stress.^5,6^ Through the folding of several hundred substrates, termed client proteins, HSP90 modulates many cellular processes beyond proteostasis including signal transduction, DNA repair and immune response that are important in several diseases such as cancer, neurodegenerative conditions, inflammation, and infectious diseases.^6–9^

Thus, HSP90 became a widely pursued drug target.^5^ HSP90 inhibitors in the clinic fall into two major lineages: 1) derivatives of natural products such as radicicol and geldanamycin; and 2) non-natural product inhibitors. The first classes of synthetic small molecule HSP90 inhibitors included purine^9^ and resorcinol derivatives (the latter sharing this motif with the natural product radicicol; Figure 1). Luminespib, ganetespib and onalespib (Figure 1) are the resorcinol derivatives that have advanced the furthest in clinical trials.^6,9,10^ Debio-0932, BIIB021 and PU-H71 are among the most clinically advanced purine derivatives.^7^ An additional class of HSP90 inhibitors harbouring a benzamide moiety has been reported, SNX-2112 being one of the most clinically advanced compounds of this class.^11^ For representative structures see Figure 1.

**Figure 1.**
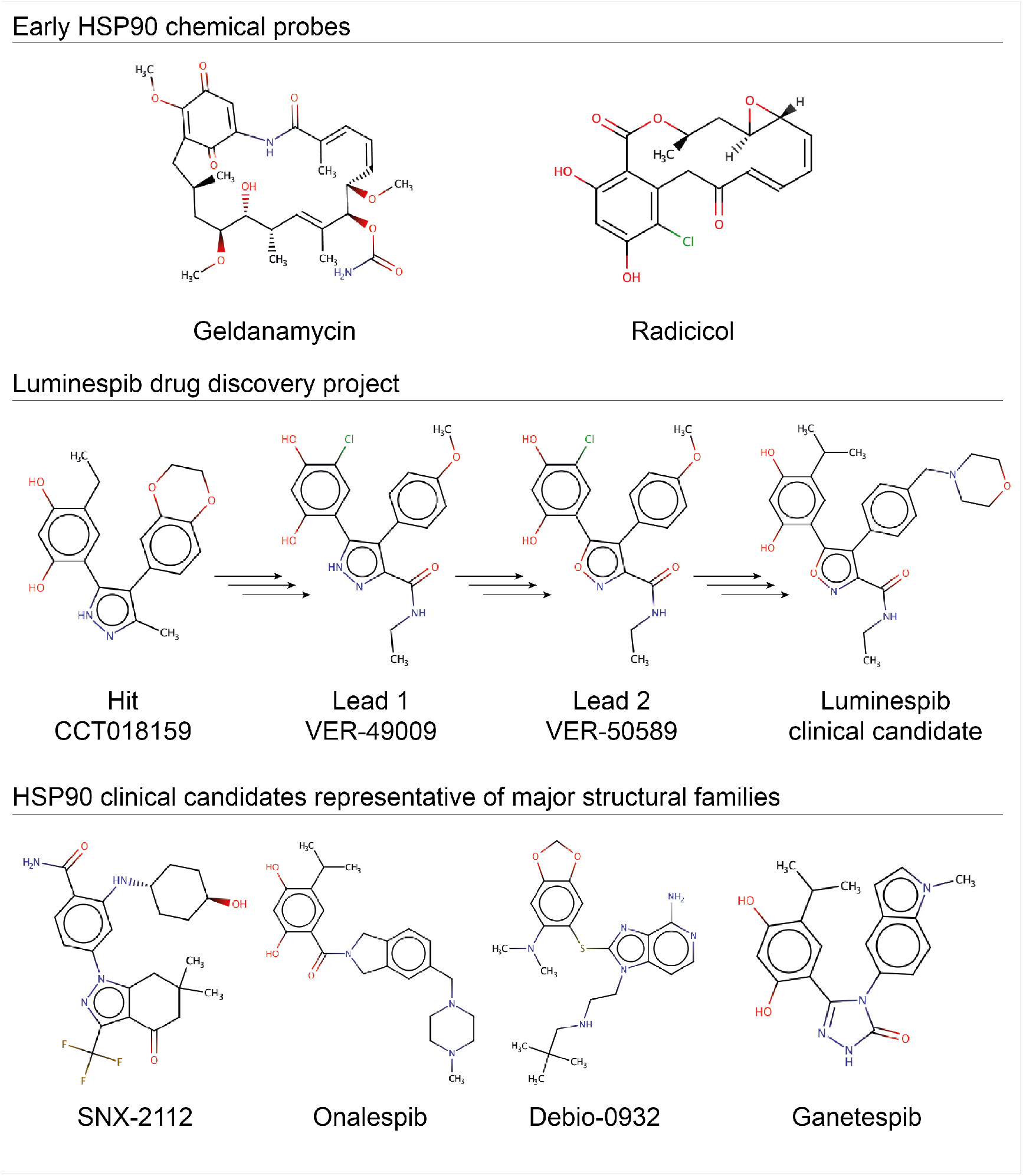
Chemical structures of selected HSP90 inhibitors.

The clinical development of HSP90 inhibitors to date has focused mainly on N-terminal ATP-site inhibitors for oncology indications. The first class of HSP90 inhibitors to be pursued clinically were geldanamycin derivatives (benzoquinone ansamycins). These provided the initial proof-of-concept and validation of HSP90 as a cancer target but generally showed modest efficacy and were limited by unfavourable properties, particularly liver toxicity, most likely due to the quinone moiety.^12,13^ Second generation synthetic HSP90 inhibitors solved some of the limitations of the first generation, including potency, solubility and hepatotoxicity, but suffered from limitations such as ocular toxicity and limited clinical activity.^14^ The most promising clinical activity was in HER2-positive breast cancer and in non-small cell lung cancer (NSCLC) patients with ALK translocations or EGFR mutations.^7^ Unfortunately, ganetespib was inactive in a Phase 3 clinical trial in combination with docetaxel in NSCLC (NCT01798485). Interestingly, the role of HSP90 in mediating drug resistance via cancer evolution is being increasingly characterized.^15–17^ Accordingly, HSP90 inhibitors could still be therapeutically relevant for the treatment of appropriately selected patient subpopulations and inclusion in drug combinations in cancer.^7^ Moreover, the potential of HSP90 inhibitors to treat other diseases remains to be comprehensively explored and recent research suggests there are exciting opportunities, such as in Alzheimer’s disease^18^ where the PU-AD has recently entered Phase 1 clinical trials (NCT03935568) or in coronavirus infections, where ganetespib (ADX-1612) has recently entered Phase 1 evaluation (https://www.aldeyra.com/pipeline-disease-areas/).

Among HSP90 clients, kinases are the most abundant protein family,^19^ many of which are themselves drug targets. Dual inhibition of HSP90 and kinases could therefore be a very attractive strategy for cancer and potentially other diseases, and a drug combinations have been suggested.^6,17,20,21^ An alternative tactic is to identify small molecules that would inhibit both HSP90 and target kinases. Of note, there is already some evidence of HSP90-kinase cross-pharmacology. The recognition of the unconventional Bergerat fold enabling ATP-binding in the GHKL family of ATPases^22^ – a family that includes both HSP90 and histidine kinases – prompted the discovery that known kinase inhibitors could inhibit DNA gyrase B^23^ and that the HSP90 inhibitor radicicol inhibited the human 3-methyl-2-oxobutanoate dehydrogenase kinase (BCKDHK, IC_50_ = 635 nM).^24^ Crystallographic evidence later showed that radicicol binds to the ATP-binding pocket of several kinases with low affinity and that the resorcinol moiety interacts via hydrogen bonding with the kinase hinge region of the bacterial sensor kinase PhoQ (kd = 715 uM)^25^ and human pyruvate dehydrogenase kinases (IC_50_ = 230 – 400 uM).^26^ A similar hinge-binding capacity has been demonstrated for other phenolic kinase inhibitor scaffolds.^27^ HSP90 inhibitor drug discovery projects, such as the one leading to the discovery of luminespib, included kinases in their off-target selectivity panels.^28,29^ However, to our knowledge, none of the clinical HSP90 inhibitors was reported to inhibit any of the kinases tested.

Several years after the publication of the initial preclinical data on luminespib, an unrelated study analysing kinase selectivity released a large biochemical screening dataset that included the original high-throughput screening (HTS) hit (CCT018159, Figure 1) and a lead compound (VER-49009, Figure 1) which led to the HSP90 inhibitor luminespib.^30^ The analysis of this dataset revealed that the hit and the lead from luminespib drug discovery project^29^ show micromolar off-target inhibition of several kinases (Supplementary Table 1).^30^ More recently, a computational analysis of pharmacological databases uncovered these and a few other dual inhibitors of HSP90 and kinases.^31^ Moreover, in the last few years four studies have rationally designed dual inhibitors of HSP90 and PDKs,^32^ BCR-ABL,^33^ or ALK,^34^ and also a triple inhibitor of HSP90, JAKs and HDAC,^35^ although the latter two of these attempts have used pharmacophore linking rather than merging.^35^ Despite the above evidence of cross-pharmacology between HSP90 and kinases, the kinase polypharmacology of HSP90 clinical candidates has not been systematically characterized and represents a very interesting area that we felt should be explored in greater depth. Moreover, the identification of off-target inhibition of kinases among the hit and lead in the luminespib project offers and unprecedented opportunity to study how kinase polypharmacology evolved across drug discovery in the absence of an explicit selection pressure.

Here, we use computational and experimental methods to systematically explore the kinase polypharmacology of representative clinical HSP90 inhibitors. We uncover the unique kinase polypharmacology of ganetespib and luminespib, the former inhibiting several kinases with nanomolar affinity. We also demonstrate that kinase polypharmacology evolved in non-obvious ways during the discovery of luminespib and we suggest new strategies to harness polypharmacology in future drug discovery campaigns.

## Results

### In silico target profiling predicts differential kinase polypharmacology between clinical HSP90 inhibitors

We used three *in silico* target profiling methods to predict computationally the protein kinase off-targets of representative non-natural product clinical HSP90 inhibitors. The three methods were: a consensus of six ligand-based chemoinformatic methods integrated in the Chemotargets CLARITY platform;^36^ the multinomial Naive Bayesian multi-category scikit-learn method available in ChEMBL;^37^ and the Similarity Ensemble Approach (SEA).^38^ All three methods use the common principle that structurally similar molecules are likely to bind to similar targets, but each one uses different similarity calculations and statistics. We selected the following representative synthetic HSP90 inhibitors from each of the chemical classes for *in silico* kinome profiling (Figure 1): three resorcinol derivatives (luminespib, ganetespib, onalespib), two purine analogs (BIIB021 and Debio-0932) and the benzamide derivative SNX-2112 (Supplementary Tables 2-4). In addition to recovering most of the known interactions between these six HSP90 inhibitors and members of the HSP90 family (Supplementary Table 1), two of the methods predicted kinases as potential off-targets of several HSP90 inhibitors that were not known (Table 1).

**Table 1.** Comparison between the number of off-target kinases predicted for HSP90 inhibitors using three *in silico* target profiling methods and the number experimentally identified by *in vitro* kinome profiling employing a radiometric catalytic inhibition assay and applying a cut-off of > 85% inhibition at 10 μM. Note that Luminespib and Ganetespib were tested in a kinome panel whilst the other HSP90 inhibitors were tested in a 16-kinase panel (Supplementary Tables).

|  |  | Method |  |  |  |
| --- | --- | --- | --- | --- | --- |
|  |  | Computational |  |  | Experimental |
| Chemical Family | HSP90 inhibitor | CLARITY<sup>36</sup> | ChEMBL<sup>63</sup> | SEA<sup>64</sup> | *In vitro* binding (KinomeSCAN®)<sup>65</sup> |
| Resorcinol derivatives | **Luminespib** | **2** | **0** | **3** | **2** |
|  | **Ganetespib** | **58** | **0** | **0** | **21** |
|  | Onalespib | 0 | 0 | 0 | 0 |
| Purine derivatives | Debio-0932 | 1 | 0 | 0 | 0 |
|  | BIIB021 | 0 | 0 | 0 | 0 |
| Benzamide derivatives | SNX-2112 | 0 | 0 | 0 | 0 |

Collectively, the methods correctly predicted most of the known interactions between the six selected HSP90 inhibitors and members of the HSP90 family (Supplementary Table 1). Additionally, two of the methods predicted protein kinases as potential off-targets of several HSP90 inhibitors that were not known (Table 1). The computational methods implemented in CLARITY predicted that ganetespib and luminespib could both inhibit kinases off-target to different extents (Table 1, Supplementary Table 2). The predictions for ganetespib had particularly high confidence (Supplementary Table 1) and, in total, CLARITY predicted that ganetespib could inhibit 56 human kinases while luminespib could inhibit only two human kinases. In contrast, onalespib, the third resorcinol derivative, was not predicted to inhibit any kinase (Supplementary Table 2). Inspection of the structure of CHEMBL156987,^37^ the compound with the highest chemical similarity to ganetespib that contributed to many of the kinase off-target predictions, showed a significant resemblance between both compounds (Figure 2). As shown in Figure 2, the maleimide ring of CHEMBL15698, likely binding to the kinase hinge region,^39^ is particularly similar to the triazolone ring of ganetespib. One carbonyl and one nitrogen atom superimpose perfectly in both heterocycles (Figure 2), which could enable ganetespib to interact with kinases. In contrast, the core oxadiazole of luminespib lacks the N-H hydrogen bond donor common to the maleimide and triazolone rings. In addition, onalespib has no five-member heterocyclic ring and its tertiary amide linker confers a more extended conformation of the molecule compared to ganetesib and CHEMBL156987 (Figure 1–2). Therefore, it is reasonable to hypothesize that the capacity of their heterocycles to interact with the hinge region of kinases and the different conformations accessible to the different scaffolds could be driving the different predictions for the four resorcinol-derived HSP90 inhibitors analysed. Interestingly, ChEMBL did not predict any kinase off-target for ganetespib, luminespib or onalespib but SEA predicted three kinases as potential off-targets of luminespib (Supplementary Tables 3-4). Overall, ganetespib and luminespib were most confidently predicted to interact with kinases off-target.

**Figure 2.**
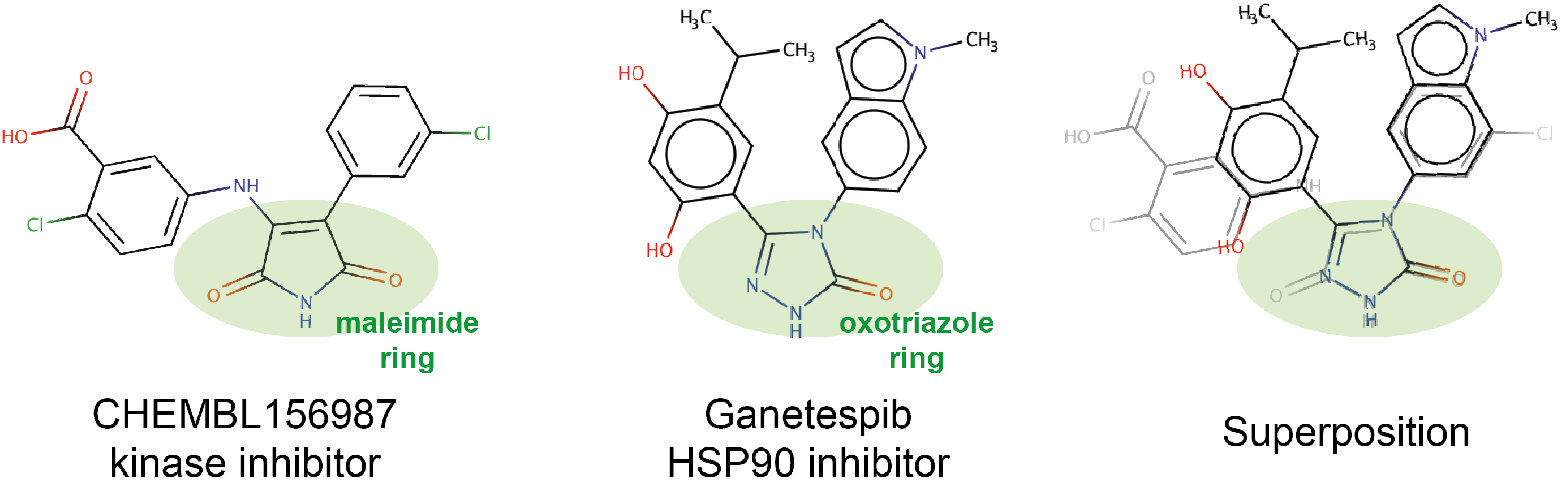
Chemical similarity between the kinase inhibitor CHEMBL156987 and ganetespib. Their similar heterocycles are highlighted in green and superimposed in the right-hand figure to highlight the complete overlap between the carbonyl and the nitrogen that are likely to interact with the kinase hinge region in CHEMBL156987.^39^

For the two purine analogs studied, CLARITY predicted only one kinase for Debio0932 (Table 1, Supplementary Table 2) and neither ChEMBL nor SEA predicted any kinase off-targets (Supplementary Tables 3-4). Thus, the computational methods used do not suggest that the purine drug candidates are likely to inhibit kinases off-target.

Finally, none of the computational methods used predicted any kinase off-targets for the HSP90 benzamide derivative SNX-2112 (Table 1, Supplementary Tables 2-4).

Overall, the computational methods we used predict that the capacity of HSP90 inhibitors to inhibit kinases off-target could vary greatly between the different chemical classes (Table 1) which could be associated particularly with the heterocyclic rings present in luminespib and ganetespib.

### In vitro kinome activity profiling uncovers differential polypharmacology between ganetespib and luminespib

In order to follow up the most promising computational predictions suggesting that luminespib and ganetespib could inhibit kinases off-target, we performed large-scale *in vitro* human protein kinome profiling of ganetespib and luminespib using Reaction Biology’s HotSpot platform (http://www.reactionbiology.com).^40^ The advantage of using this radiometric assay is that it measures inhibition of catalytic activity as opposed to the binding assays employed by other large-scale kinome profiling platforms that can report binding that does not translate into inhibition of catalytic activity.^41^ At the time of performing the experiments, the kinome panel used was the largest commercially available platform measuring catalytic activity and comprised 583 assays corresponding to 382 unique human kinases (74% of the human protein kinome).^42^ Additionally, 174 assays included mutated forms of kinases, 17 translocated products and 10 other genomic aberrations (Supplementary Table 5). The kinome profiling was performed at 10 μM concentration to identify both low and high potency kinase off-targets.

The results of the *in vitro* kinome profiling validated our computational prediction that ganetespib and luminespib inhibit kinases off-target while uncovering significant differences between these two clinical HSP90 inhibitors. As illustrated in Figure 3,^43^ ganetespib inhibited 20 native kinases and the fusion TRKA-TFG by more than 85% at 10 μM whereas luminespib inhibited only two native kinases (Supplementary Table 5). Both drugs also inhibited several mutated forms of kinases. Given the variability associated to single-point high-concentration screens, we selected an 85% cut-off to increase the likelihood of the IC_50_ being lower than the tested concentration. Interestingly, the two kinases inhibited by luminespib, ABL1 and ABL2, were also inhibited by ganetespib with a higher percentage of inhibition at 10 μM. As illustrated in Figure 3, the kinase off-target activities of ganetespib are relatively widely distributed across the kinome tree, while the two kinases that luminespib was found to inhibit belong to the Tyrosine Kinase (TK) group.

**Figure 3.**
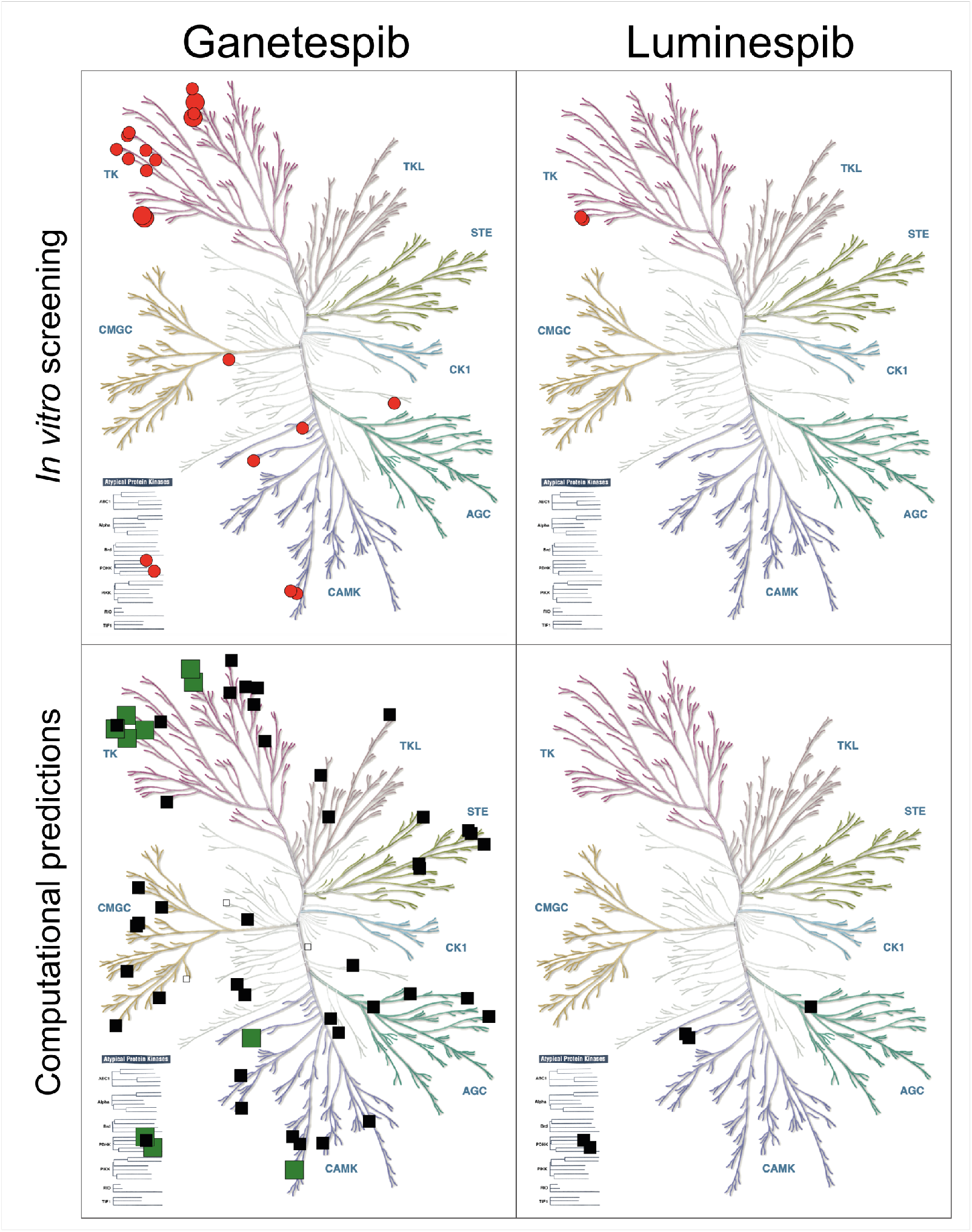
Experimental and computational results superimposed onto kinome trees for the HSP90 clinical inhibitors ganetespib (left panel) and luminespib (right panel) using KinMap.^43^ The top panel displays the experimental hits of the *in vitro* kinome screening using the catalytic inhibition assay which are represented as red dots (see Methods). Large red dots represent submicromolar interactions (IC_50_ < 1 μM) while medium red dots represent micromolar interactions either calculated in concentration-response experiments or expected based on the screening results (% inhibition at 10 μM > 85% and % inhibition at 1 μM < 85%). The bottom panel merges the *in silico* predictions of the three computational methods used (see Results and Methods) distinguishing between true positives (large green squares), false positives (medium black squares) and kinases which could not be experimentally validated (small white squares).

Of the 58 human kinases that CLARITY predicted ganetespib or luminespib could inhibit, 10 were correctly predicted whilst 45 were false positives and 3 were not available in the selected kinase panel (Figure 3, Table 1 and Supplementary Table 2). Therefore, CLARITY had a sensitivity (recall) of 0.48 (10/21) regarding the kinase off-targets of ganetespib that showed >85% kinase inhibition *in vitro* at 10 μM whilst its precision was 0.17 (10/58). ChEMBL did not predict any kinase off-targets for luminespib or ganetespib and SEA predicted three kinases as potential off-targets of luminespib, none of which was validated *in vitro*. Therefore, the recall and precision was 0 for both methods. (Supplementary Table 3-4). Overall, the sensitivity and precision between these three computational methods was significantly different, despite their sharing of the same underlying principles. CLARITY’s integration of several methods appears to be advantageous in this particular example. It would be important to thoroughly study the dependencies of all these methods to the different fingerprints and statistical approaches used and whether a better integration of methods could improve the sensitivity and precision of *in silico* polypharmacology prediction further.

### Concentration-response determinations confirm ABL1, ABL2, DDR1 and TRKA-TFG as submicromolar off-targets of ganetespib

For further follow-up, we performed a secondary screening round at 1 μM using the same radiometric assay. We prioritised the 20 native kinases inhibited by more than 85% by ganetespib at 10 μM (Supplementary Table 5) and also included the TRKA-TFG (TRK-T3) fusion. From these, only ABL2 and DDR1 were found to be inhibited by ganetespib by more than 85% at 1 μM (Figure 3, Supplementary Table 5). Interestingly, both belong to the TK family (Figure 3).^43^

Next, we validated the most potent interactions using the radiometric 10-point concentration-response inhibition assay in triplicate (Supplementary Table 6). For luminespib, we selected the two native kinases whose activity was inhibited by more than 85% at 10 μM, namely, ABL1 and ABL2 (Supplementary Table 5). For ganetespib we selected ABL2 and DDR1, both inhibited by more than 85% at 1 μM. We also tested ganetespib against ABL1 and TRKA-TFG because of their potential therapeutic relevance, and since both of them inhibited by more than 80% at 1 μM (Supplementary Table 5). As can be observed in Figure 4, ganetespib exhibits submicromolar IC_50_ values for all of the four kinases tested, the most sensitive of which is ABL2 (IC_50_ = 215 nM). In contrast, luminespib exhibits micromolar IC_50_ values for the two kinases tested, the most sensitive of which is ABL1 (IC_50_ = 3,391 nM).

**Figure 4.**
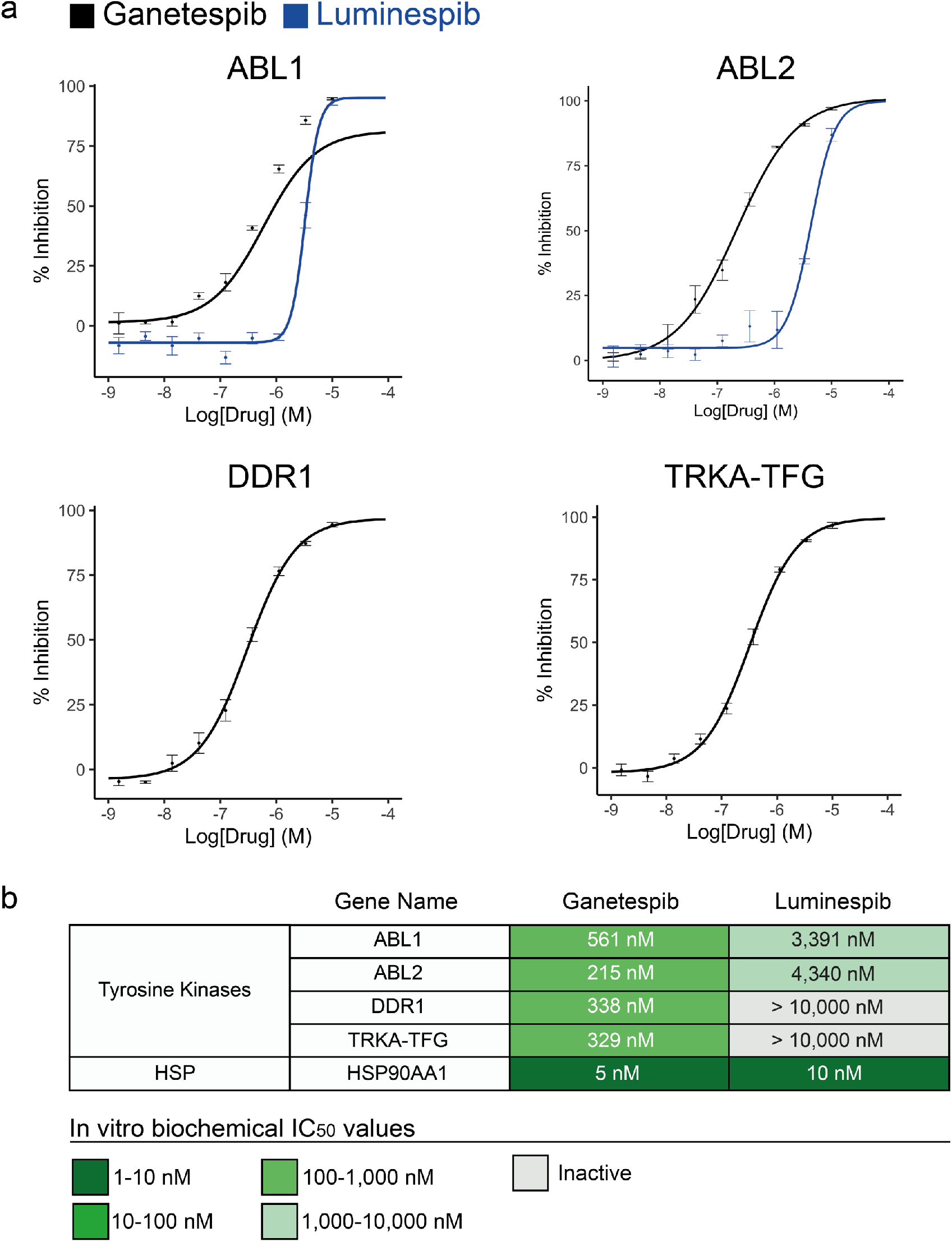
Concentration-response curves and IC_50_ values (n=3) of the most potent kinase off-target interactions of luminespib and ganetespib. **a** concentration-response curves of the interactions of ganetespib and luminespib with ABL1 and ABL2 (top) and ganetespib with DDR1 and TRKA-TFG (bottom). **b** Table summarising the calculated IC_50_ values for the kinase off-targets of ganetespib and luminespib. HSP90AA1 IC_50_ average values obtained from ChEMBL are also included for comparison (see also Supplementary Table 1).^37^

Overall, the different kinase polypharmacology profiles of these two HSP90 inhibitors has been experimentally confirmed, with ganetespib showing far greater kinase polypharmacology and more than 10-fold increased affinity for kinases than luminespib, which displays only very modest micromolar affinity for only two kinases. To our knowledge, this is the first report of submicromolar kinase off-targets of ganetespib.

### Kinase polypharmacology is not ubiquitous among HSP90 inhibitors

Having systematically characterized the kinase polypharmacology of ganetespib and luminespib, we wished to determine if this is a general property of all HSP90 inhibitors. Accordingly, we selected five additional representative HSP90 inhibitors, at least one from each chemical class, including both non-natural product inhibitors and natural products. Thus, we chose the natural products geldanamycin and radicicol, the resorcinol derivative onalespib, the purine derivative Debio-0932 and the benzamide SNX-2112, the latter three being also clinical candidates (Supplementary Table 7). These five HSP90 inhibitors were screened at 10 μM concentration against the 15 kinases with >50% inhibition at 1 μM of ganetespib). The final 16-kinase panel also included LYN B as the kinase with the highest inhibition by luminespib that is not inhibited by ganetespib. Interestingly, none of the additional HSP90 inhibitors tested displayed significant activity against any of the 16 kinases tested (Supplementary Table 7). Thus, it appears that off-target kinase pharmacology seen with ganetespib and, to a lesser extent, luminespib, is not a general property of HSP90 inhibitors.

### Docking experiments to study HSP90-kinase cross-pharmacology at the atomic level

In order to study whether the unique capacity of HSP90 clinical inhibitors ganetespib and luminespib to inhibit kinases off-target was facilitated by their heterocyclic rings, we used molecular docking. We selected ABL1 as a representative kinase because it is the most potent off-target kinase for both inhibitors. We used the structure of ABL1 cocrystalized with tetrahydrostaurosporine, because tetrahydrostaurosporine is the inhibitor co-crystalized with ABL1 most similar to ganetespib and its closest kinase inhibitor CHEMBL156987 – which had enabled the off-target kinase prediction of ganetespib (Figure 2, Methods). We subsequently docked the five selected HSP90 inhibitors that were demonstrated not to inhibit ABL1 (Figure 1, Supplementary Table 7) as well as ganetespib and luminespib. All docking were performed using MOE (Methods).

The docking results were consistent with our hypothesis that the triazolone ring of ganetespib would be able to interact with the kinase hinge region, thus providing an explanation for the highest affinity of ganetespib for a larger number of kinases when compared to luminespib and the rest of HSP90 inhibitors studied. As illustrated in Figure 5, the interaction maps derived from the best docking poses for ganetespib, luminespib and onalespib show how ganetespib is predicted to interact with the kinase hinge region, reproducing the double hydrogen bonding pattern of tetrahydrostaurosporine. Both tetrahydrostaurosporine and ganetespib are predicted to interact via hydrogen bonds with the ABL1 hinge region residues Met 318 and Glu 316. Luminespib’s best binding pose predicted by docking places the isoxazole in a different area of the binding site, although one of the hydroxyl groups of the resorcinol ring would still be able to form one hydrogen bond with the hinge residue Met 318. This same interaction of the resorcinol ring with Met 318 is maintained in radicicol’s best pose (Supplementary Figure 4) and is similar to the published crystal structures of radicicol with other kinases.^25,26^ Onalespib, the third resorcinol derivative, is not predicted to make any hydrogen bond with the hinge region of ABL1. Thus, the resorcinol moiety, albeit that it has been shown to interact with the kinase hinge region, is not sufficient to achieve potent inhibition as illustrated by radicicol and onalespib – both inactive in the kinase panel tested. None of the rest of the HSP90 inhibitors from other chemical families that we found to be inactive against the 16-kinase panel (Supplementary Table 7) were predicted to mimic the double hydrogen bond of tetrahydrostaurosporine with residues Met 318 and Glu 316 (Supplementary Figures 1-4). However, the best docking poses predicted that some of the above inhibitors would be able to make hydrogen bonds with hinge residues. The docking scores were poor predictors of ABL1 inhibition activity, in line with the ample evidence that docking performs better at predicting the binding pose of known inhibitors than at predicting binding affinity.^44^ Overall, our docking results support our hypothesis that the triazolone ring of ganetespib and its capacity to interact with the kinase hinge region could be responsible for driving its kinase polypharmacology.

**Figure 5.**
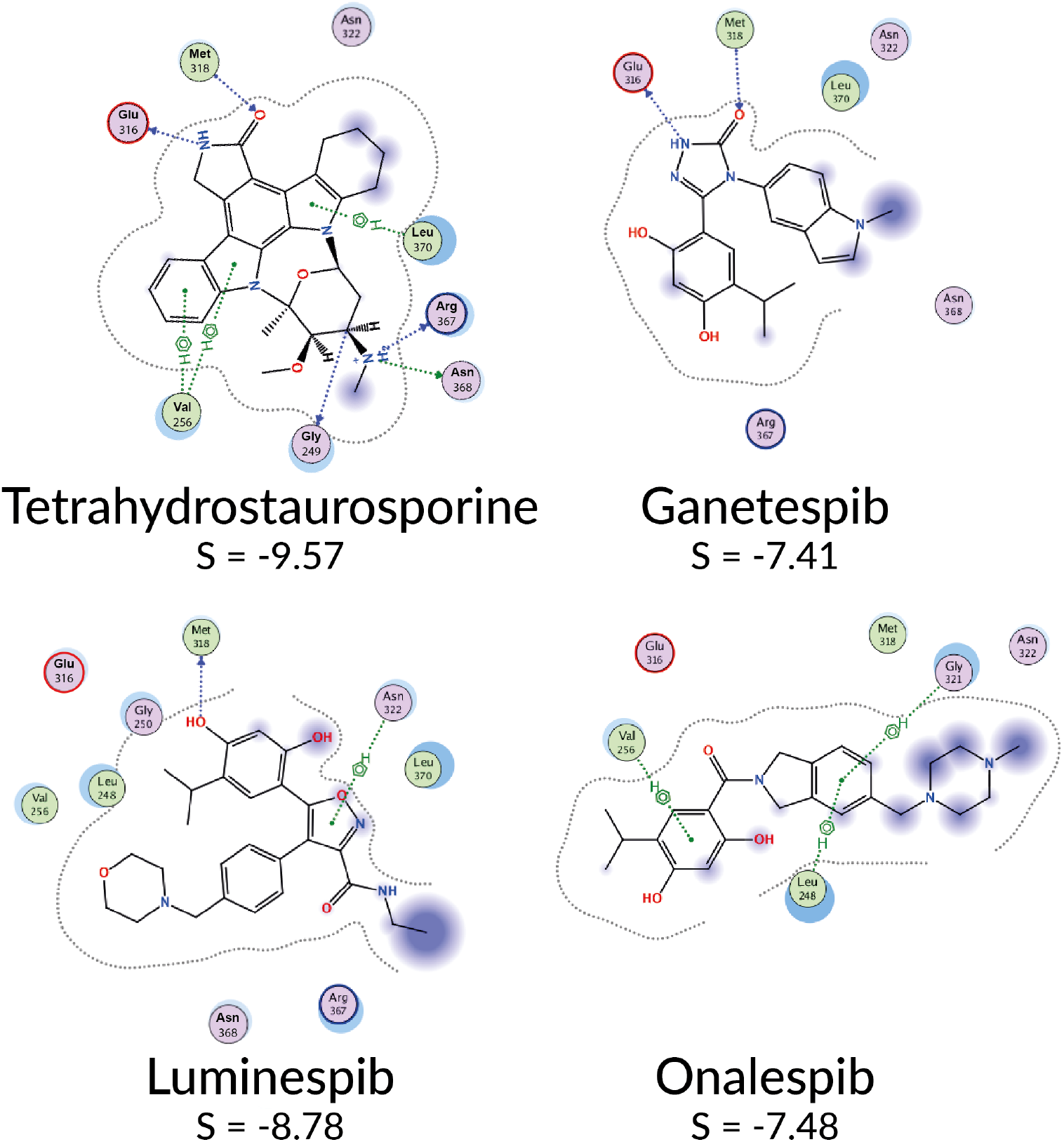
Analysis of ABL1-ligand interactions of different HSP90 inhibitors. The cocrystalized ABL1 inhibitor tetrahydrostaurosporin and three clinical HSP90 inhibitors of the resorcinol family (ganetespib, luminespib and onalespib) were docked using MOE and the best docking pose according to MOE score was analysed using MOE’s ligand interactions tool. The interaction diagrams for each of the four inhibitors are displayed highlighting only the protein residues interacting with at least one of the inhibitors. Ganetespib is the only HSP90 inhibitor predicted to mimic the double hydrogen bond that tetrahydrostaurosporin dispays for interaction with the kinase hinge region.

### Evolution of kinase polypharmacology across luminespib drug discovery

Luminespib was discovered by our academic drug discovery unit in collaboration with Vernalis.^28,29^ The chemical structures of the initial 3,4-diarylpyrazole resorcinol screening hit CCT018159,^45,46^ advanced lead compounds VER-49009^47^ and VER-50589^48^, and drug candidate itself (Figure 1) were disclosed and the compounds are commercially available. This offers an opportunity to study how kinase polypharmacology evolved across this drug discovery project in the absence of any explicit selection for kinase activity. Accordingly, we screened the hit and the two leads for kinase off-target inhibition using the 16-kinase panel and the same *in vitro* catalytic assay used previously. This panel includes nine of the 14 kinase off-targets of luminespib inhibited by more than 50% at 10 μM and also the most potent off-targets of ganetespib that are not inhibited by luminespib (e.g. TRKA) in order to sample both off-targets that could be shared and those that could be different between luminespib and its precursor hit and lead compounds. Indeed, the screening did uncover off-targets that are in common and those that are different between the four compounds (Supplementary Table 7).

As can be observed in Figure 6, polypharmacology evolved across luminespib’s drug discovery history. The 4,5-diarylpyrazole screening hit CCT018159 and the final clinical candidate luminespib have more off-targets than the intermediate lead compounds VER-49009 and VER-50589 in the kinase panel used. Introduction of the 3-carboxamide substituent into the early lead 4,5-diaryl 3-carboxamido pyrazole VER-49009 significantly increased the off-target activity for PDK2/PDK4, which was maintained in subsequent HSP90 inhibitor optimization. Other kinase inhibitory activities of the hit were reduced by introduction of the 3-carboxamide substituent, itself an important contributor to affinity for HSP90. The change of five-membered heterocycle from the 4,5-diaryl 3-carboxamido pyrazole VER-49009 to the 4,5-diaryl 3-carboxamido isoxazoles VER-50589 and luminespib was associated with a large reduction of the TRKA off-target activity observed for the pyrazoles. The principal structural differences between the 4,5-diaryl 3-carboxamido isoxazoles VER-50589 and luminespib are the introduction of a basic morpholino substituent and the change of the resorcinol chloro-substituent to the larger isopropyl group, while the core isoxazole amide scaffold is preserved. These changes in peripheral substitution are associated with an increase in the number of off-target kinases inhibited. The differential dependence of the off-target kinases on the presence of different functional groups suggests that in some cases the scaffolds could adopt different binding poses between kinases, driven by different key interactions. Crystallographic determination of binding modes or more extensive structure-activity relationships for the off-target activities are required to investigate this possibility. All of the compounds have different polypharmacology profiles that evolved in non-obvious ways. For example, the hit CCT018159 and luminespib share >70% inhibition of PEAK1 that is not inhibited by either of the intermediate lead compounds (< 24% inhibition). The hit CCT018159 inhibits DDR1 (81% inhibition), a kinase that is inhibited by only 29% or less by the other compounds. Perhaps surprisingly, luminespib has a more similar off-target profile to the hit CCT018159 than to the lead compounds VER-49009 and VER-50589 and has four unique off-targets not shared with any of the hit and lead compounds. Overall, polypharmacology can evolve non-linearly and is not necessarily present in the hit or lead compounds.

**Figure 6.**
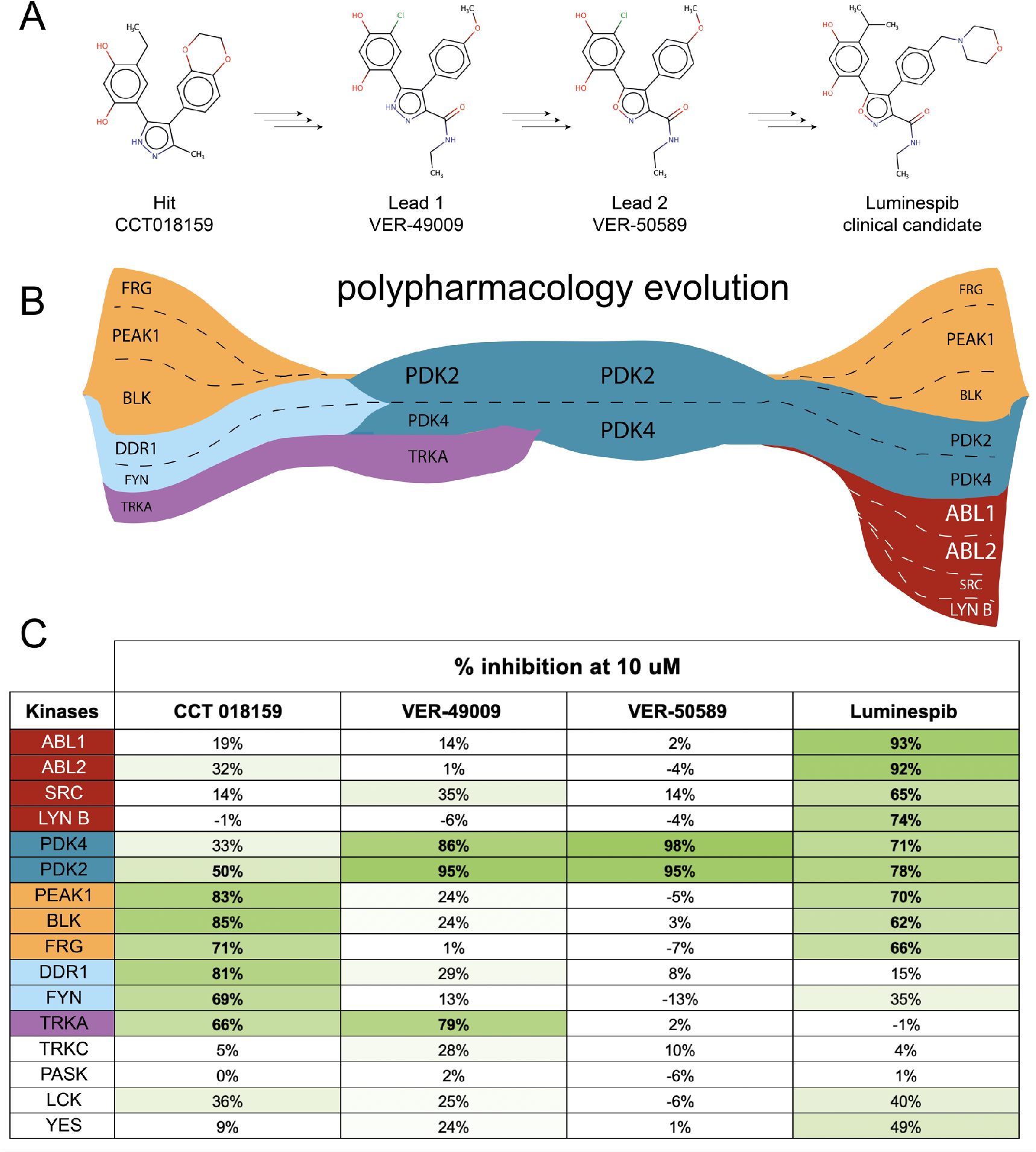
Change in kinase polypharmacology during the evolution of HSP90 inhibitors resulting in the discovery of luminespib. A. the chemical structures of the screening hit, the two lead compounds and the clinical candidate luminespib are displayed. B. we plot the evolution trajectories of the micromolar kinase off-targets of each of these compounds below the corresponding chemical structure considering a >50% inhibition at 10 μM cut-off (Supplementary Table 7). C. Since the consideration of a single cut-off value can be inaccurate, on the bottom panel we display the median % inhibition values (n=2) for each compound-kinase pair (green color intensity is proportional to % inhibition) to trace how kinase polypharmacology evolved across this particular drug discovery project.

## Discussion

In this study, we have performed a comprehensive computational and experimental characterization of the protein kinase polypharmacology of clinical HSP90 inhibitors. We have demonstrated that, in line with our computational predictions, off-target kinase pharmacology is a property of the clinical candidate ganetespib and, to a lesser extent, luminespib, another resorcinol clinical candidate. In contrast, this off-target kinase profile is not an inherent property of all HSP90 inhibitors as the natural products geldanamycin and radicicol and the non-natural product clinical candidates Debio-0932, onalespib and SNX-2112 were inactive against the 16-kinase panel used (Figure 3, Supplementary Table 7). Ganetespib inhibits 21 native kinases *in vitro* with micromolar affinity, 4 of them with affinities below 1 μM. In contrast, luminespib inhibits only 2 kinases with micromolar affinity (Figure 4). Of interest, from the 21 off-target kinases, 16 have been reported to be HSP90 client proteins, further emphasizing the potential relevance of their off-target inhibition (Supplementary Table 5). It is also worth noting that kinase inhibitors can lead to enhanced degradation, and therefore reduced expression of the target kinase, by blocking recruitment of kinases to HSP90 by CDC37.^49^ It is interesting to speculate that this effect could have indirectly favoured the selection of dual HSP90-kinase inhibitors in cellular assays that measure client protein depletion and cell growth inhibiton. Therefore, it is possible that the kinase polypharmacology of ganetespib and luminespib could have clinical relevance. Moreover, these results might unlock new opportunities for the rational design of dual HSP90-kinase inhibitors with improved therapeutic properties and provide new lessons for better harnessing polypharmacology in drug discovery.

Of the three computational methods used, two of these correctly predicted different kinases as off-targets of ganetespib and luminespib despite there being significant differences in the sensitivity and precision of each method (Figure 3, Supplementary Tables 2 - 4). As we have recently shown in the case of PARP inhibitors,^50^ the methods differ in specific results and may not predict the exact kinase off-target, but collectively they correctly predict target classes and thus can usefully guide further experimentation. Despite these differences, when used together, the methods were able to anticipate the kinase polypharmacology of ganetespib and luminespib as compared to other HSP90 inhibitors (Figure 3). Accordingly, we recommend the use of these computational methods in conjunction to maximize recovery when predicting for potential off-targets.

Despite the clear usefulness of computational methods, experimental family-wide biochemical profiling using a radiometric catalytic assay was able to identify a significant number of additional protein kinase off-targets *in vitro* that had not been predicted *in silico* (Figure 3). This indicates the importance of investigating polypharmacology effects across a particular target family as a means to increase the number of identified off-targets. Note, however, that current kinome panels do not yet provide complete family coverage. It is therefore important that these panels continue to expand to facilitate the identification of a larger number of kinase off-targets in the future.

Understanding ganetespib’s polypharmacology at the atomic level could be particularly important to facilitate the rational development of new HSP90-kinase multitarget inhibitors. Given the dual role of HSP90 in folding many kinases and in mediating drug resistance,^15,19,51^ simultaneous inhibition of HSP90 and therapeutically important kinases could be an interesting strategy to prevent or delay the persistent problem of cancer drug resistance.^15,20,52,53^ There is a high chemical similarity between ganetespib and its closest kinase inhibitor, CHEMBL156987. Both inhibitors have similar heterocyclic rings (Figure 2), which in the case of CHEMBL156987 was suspected by the researchers that discovered this small molecule to bind to the kinase hinge region of GSK3.^39^ Given the importance of the hinge region for kinase binding, we used docking to investigate whether the triazolone ring of ganetespib could be binding to the kinase hinge region. Indeed, docking results supported this hypothesis as the triazolone appears to be capable of reproducing the double hydrogen bond with two hinge residues observed in the cocrystal of tetrahydroxystaurosporin with ABL1 (Figure 5). Although the resorcinol moiety of radicicol has been shown in two crystal structures to interact with the kinase hinge region of PhoQ and pyruvate dehydrogenase kinases and confer modest binding,^25,26^ the resorcinol derivatives radicicol and onalespib were found to be inactive (<22% inhibition @ 10 μM) in the kinase panel tested, strengthening the hypothesis that the triazolone is the key moiety enabling the kinase polypharmacology of ganetespib. The best docking poses of the other HSP90 inhibitors studied here were not capable of reproducing this double hydrogen bond with ABL1 hinge region residues (Figure 5, Supplementary Figures 1 - 4). However, it would be important to confirm these docking predictions with x-ray crystallography. Overall, the triazolone ring of ganetespib emerges as a potential privileged structure for simultaneous inhibition of HSP90 and kinases and an ideal starting point for future rational multi-target drug design endeavours.

We propose that the potent nanomolar off-target activities of ganetespib for several kinases could be therapeutically relevant and thus should be investigated further, particularly regarding the potential clinical impact of inhibiting ABL1, ABL2, DDR1 and TRKA-TFG. There are no data in the public domain on the profiling of ganetespib in large-scale cancer cell line panels such as the Sanger’s Genomics of Drug Sensitivity in Cancer^54^ or the Broad’s Cancer Therapeutics Response Portal^55^ but available evidence^56^ suggests that it could have different cellular effects when compared to other HSP90 inhibitors. Of interest, a recent quantitative proteomic study in a human bladder cancer cell line, identified differences in protein expression changes induced by ganetespib (0.5 - 1 μM) versus luminespib (0.1 – 0.25 μM).^56^ Although the full dataset was not made publicly available, data for some of the unique off-targets we identified for ganetespib like PLK1 (Supplementary Table 5) were disclosed and revealed differential expression following treatment with luminespib (2-fold decreased) or ganetespib (5-fold decreased).^56^ Thus, the observed differences in expression may reflect the differential polypharmacology we observed between these clinical candidates.

It would be important to investigate whether the nanomolar affinity for kinases identified in the present work and the observed differences in expression between luminespib and ganetespib at the cellular level^56^ have clinical implications. HSP90As are abundant and ubiquitous proteins and overexpressed under stress conditions such as cancer.^5^ Moreover, ganetespib displays at least 40-fold selectivity for HSP90 family members over kinases (Figure 3). Despite, to our knowledge, the unbound concentration of ganetespib in human plasma has not been reported, the micromolar total plasma Cmax values of ganetespib achieved in clinical studies (Cmax, total = 6 - 12 μM)^57^ could allow drug concentrations compatible with inhibition of potent off-target kinases to be reached in patients. This may potentially be limited due to competition from HSP90 family members. In particular, it would be important to clarify whether ganetespib can inhibit kinases in humans. For example, pharmacodynamic biomarkers of ABL1 inhibition could be tested retrospectively in biosamples from patients that had been treated with ganetespib (which inhibits ALB1) and compared to results for patients treated with onalespib (which does not inhibit ABL1) in order to demonstrate whether ABL1 was engaged by ganetespib at clinically achieved concentrations.

Ganetespib has failed to show overall benefit in non-stratified clinical trials of NSCLC. However, some of the cancer types where ganetespib has been tested could be stratified retrospectively for aberrations in the kinase off-targets identified in this work. For example, TRKA translocations affect <1% of NSCLC, a disease in which ganetespib showed a disappointing response overall.^58^ Given the enormous economic costs of late-stage clinical trial failures such as ganetespib in NSCLC,^59^ any exceptional patient responders in these trials could be evaluated retrospectively using patient selection biomarkers for kinase off-targets of ganetespib such as TRKA. However, it would be important to first validate that TRKA is engaged by ganetespib in cellular and animal models. If TRKA target engagement was validated in such preclinical models, it may be also worth exploring the repurposing of ganetespib in cancer subtypes not yet investigated where kinase off-targets are both validated cancer drivers as well as HSP90 clients. For example, TRKA has been shown to be an HSP90 client^19^ and paediatric TRKA-translocated cancers have shown strong responses to the TRKA kinase inhibitor larotrectinib, but suffer from the inevitable appearance of drug resistance.^60^ In this context, it would be particularly interesting to explore whether the use of ganetespib, or more potent, newly-designed dual TRKA-HSP90 inhibitors, could delay or prevent the appearance of drug resistance in TRKA-translocated cancers. Overall, our results may unlock new opportunities to harness polypharmacology for a more precise and personalized development of ganetespib in selected cancer subpopulations.

Today, there is a growing interest in the rational design of multi-target drugs.^3,61^ However, it is still very challenging to identify targets that can be simultaneously inhibited with a single compound and that are both therapeutically relevant for a specific disease.^3^ Here, we have benefited from the commercial availability of the screening hit, advanced lead compounds and clinical candidate to study how polypharmacology evolved during luminespib’s drug discovery history. Interestingly, no explicit selection pressure was applied as the kinase polypharmacology of these compounds was discovered *a posteriori*. To our surprise, both the clinical candidate luminespib and the screening hit CCT018159 display greater kinase polypharmacology than the intermediate lead compounds VER-49009 and VER-50589 (Figure 6, Supplementary Table 7). This observation might be an important consideration when using lead compounds for proof-of-concept experiments. Although the 16-kinase panel used here is limited, we show how polypharmacology can evolved during a drug discovery project. All the compounds studied display a unique kinase polypharmacology and the molecular target profile of the drug candidate is not completely present in the screening hit or lead compounds (Figure 6). Albeit this is one single example, our findings suggest that many potentially interesting off-targets could be missed if only the clinical candidate is comprehensively profiled. Different kinase selectivity trajectories have also been observed during fragment growing of ATP-competitive kinase inhibitors, where minimal changes in the fragment can lead to new interactions with the target(s) that alter the kinase selectivity pattern of the fragment.^62^ We propose that a more systematic exploration of off-target pharmacology earlier in drug discovery campaigns could be helpful in the identification of multi-target drug design opportunities, or steering polypharmacology towards more advantageous outcomes. Multi-target chemical series could potentially be developed in parallel with chemical series aiming at single target inhibition and tested in follow-up phenotypic or efficacy experiments. For example, in the case of luminespib a dual HSP90-TRKA series inhibitor could potentially have been developed in parallel with a series aiming to design out any off-target kinase pharmacology. We also propose that computational prediction of polypharmacology, despite its limitations, might be very valuable at the earlier stages of drug discovery where comprehensive experimental profiling may not be cost-effective. Overall, the characterization of polypharmacology at earlier stages of drug discovery may unlock new multi-target drug design opportunities.

In summary, our study demonstrates the unique kinase polypharmacology of the HSP90 inhibitors ganetespib and luminespib. In particular, ganetespib inhibits several human protein kinases with nanomolar affinity, a number of them being HSP90 clients. If proven to be therapeutically relevant, this could open new avenues for the retrospective identification of better patient selection biomarkers, and new opportunities for drug development that reduce the enormous societal cost of discontinued clinical candidates. Moreover, they could also facilitate the repurposing of ganetespib for new precision oncology indications or contribute to explain differences in side-effects between these drugs. Finally, our results demonstrate that polypharmacology can evolve non-linearly in drug discovery and we recommend the computational and experimental characterization of polypharmacology earlier in drug discovery projects to harness untapped opportunities for multi-target drug design.

## Acknowledgements

A.A.A. is primarily supported by a Wellcome Trust Sir Henry Wellcome Postdoctoral Fellowship (204735/Z/16/Z); the People Programme (Marie Curie Actions) of the 7th Framework Programme of the European Union (FP7/2007–2013) under REA grant agreement no. 600388 (TECNIOspring programme) and the Agency of Business Competitiveness of the Government of Catalonia, ACCIO. B.A.-L., P.C., I.C. and P.W. are funded a Cancer Research CRUK programme grant to the Cancer Research UK Cancer Therapeutics Unit (grant C309/A11566), a Wellcome Trust biomedical resource and technology grant to develop the Chemical Probes Portal (212969/Z/18/Z) and a Cancer Research UK (CRUK) Drug Discovery Committee strategic award to develop canSAR (C35696/A23187). P.W. is a CRUK Life Fellow. P.W. and B.A.L. are members of the CRUK ICR/ Imperial Convergence Science Centre. We acknowledge support from CRUK and the NHS funding to the CRUK Centre and Biomedical Research Centre respectively at the ICR and Royal Marsden NHS Foundation Trust. The authors thank many colleagues and collaborators for discussions and valuable input into the preparation of this manuscript. In particular, the authors thank Udai Banerji for valuable discussions and advice.

## Author contributions

A.A.A., P.A.C., P.W., and B.A.-L. designed the research. A.A.A. performed the computational and managed the experimental work. A.A.A., P.A.C., I.C., P.W., and B.A.-L. conducted data analysis and interpretation. A.A.A., P.A.C, I.C., P.W., and B.A.-L. contributed to manuscript preparation.

## METHODS

### In silico target profiling

The chemical structures of the HSP90 inhibitors were downloaded from ChEMBL (canonical SMILES).^63^ Three different *in silico* methods were used to predict the kinase off-targets of selected HSP90 clinical candidates, all exploiting the chemical similarity principle. The first method used was CLARITY (https://www.chemotargets.com), which computes a predefined consensus of six ligand-based chemoinformatic methods.^36^ The second method employed was the Similarity Ensemble Approach (SEA) (http://sea.bkslab.org/) with default parameters.^38^ And finally, the third method selected was the similarity-based method available through the ChEMBL website (https://www.ebi.ac.uk/chembl/).^63^ Supplementary Tables 2 - 4 list the predictions obtained from these three computational methods.

### In vitro kinase radiometric assays

Reaction Biology’s HotSpot platform (http://www.reactionbiology.com)^40^ was used at 10 μM for kinome profiling and at 1 μM and/or 10-point concentration-response to validate the most potent hits. The radioisotope binding assay was designed to directly detect the true product without the use of modified substrates, coupling enzymes, or detection antibodies thus enabling to directly measure inhibition of catalytic activity. Test or control compounds were incubated with kinase, substrate, cofactors, and radioisotope-labeled ATP (32P-ɣ-ATP or 33P-ɣ-ATP). The reaction mixtures were then spotted onto filter papers, which bind the radioisotope-labeled catalytic product. Unreacted phosphate is removed via washing of the filter papers.^41^

### Docking experiments

From the PDB, we selected the crystal structure corresponding to the wild type kinase domain ABL1 and co-crystalized with the most similar small molecule to ganetespib and its closest kinase inhibitor CHEMBL156987 that enabled the kinase off-target predictions (PDB ID: 2HZ4, Ligand: tetrahydrostaurosporine (4ST)). The PDB file was prepared using the standard preparation method QuickPrep implemented in MOE 2018.01 (https://www.chemcomp.com/Products.htm). The mol files of the seven HSP90 inhibitors docked were downloaded from ChEMBL. The binding site was described using the cocrystalised ligand and docking was performed using standard variables for rigid docking. Best docking poses were analysed with MOE’s Ligand Interactions tool.

