## Supplementary Figures for "Evolution of kinase polypharmacology across HSP90 drug discovery"

| Supplementary | Title | Page # |
|---|---|---|
| Supplementary Figure 1 | Protein-ligand interactions scheme for the docking pose with the top MOE score of the clinical HSP90 inhibitor SNX-2112 in ABL1 kinase. | 3 |
| Supplementary Figure 2 | Protein-ligand interactions scheme for the docking pose with the top MOE score of the clinical HSP90 inhibitor Debio-0932 in ABL1 kinase. | 4 |
| Supplementary Figure 3 | Protein-ligand interactions scheme for the docking pose with the top MOE score of the clinical HSP90 inhibitor geldanamycin in ABL1 kinase. | 5 |
| Supplementary Figure 4 | Protein-ligand interactions scheme for the docking pose with the top MOE score of the clinical HSP90 inhibitor radicicol in ABL1 kinase. | 6 |

**Supplementary Figure 1.** Docking pose with the top MOE score (S = -8.20) for SNX-2112 in ABL1 kinase. The MOE ligand interaction tool was used to generate the schematic diagram of protein-ligand interactions.

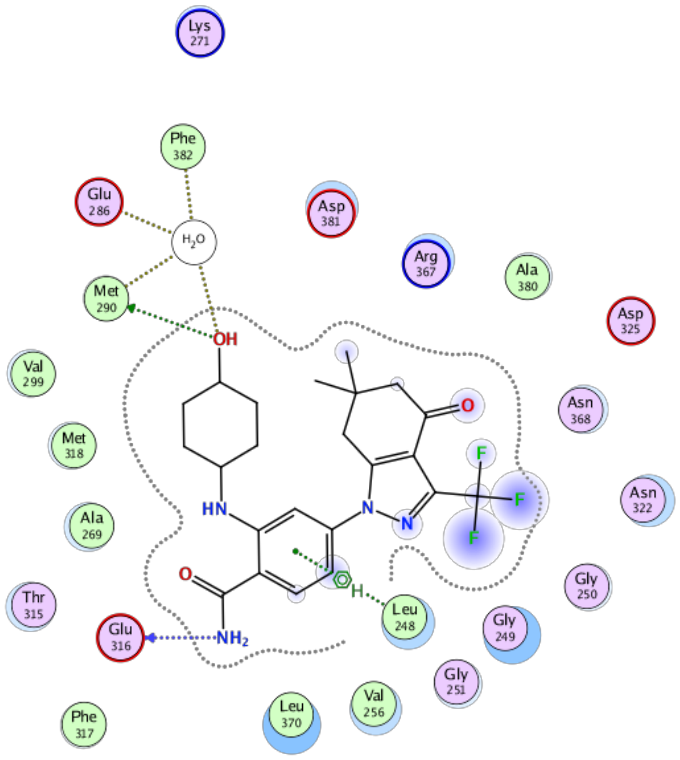

**Supplementary Figure 2.** Docking pose with the top MOE score (S = -8.02) for Debio-0932 in ABL1 kinase. The MOE ligand interaction tool was used to generate the schematic diagram of protein-ligand interactions.

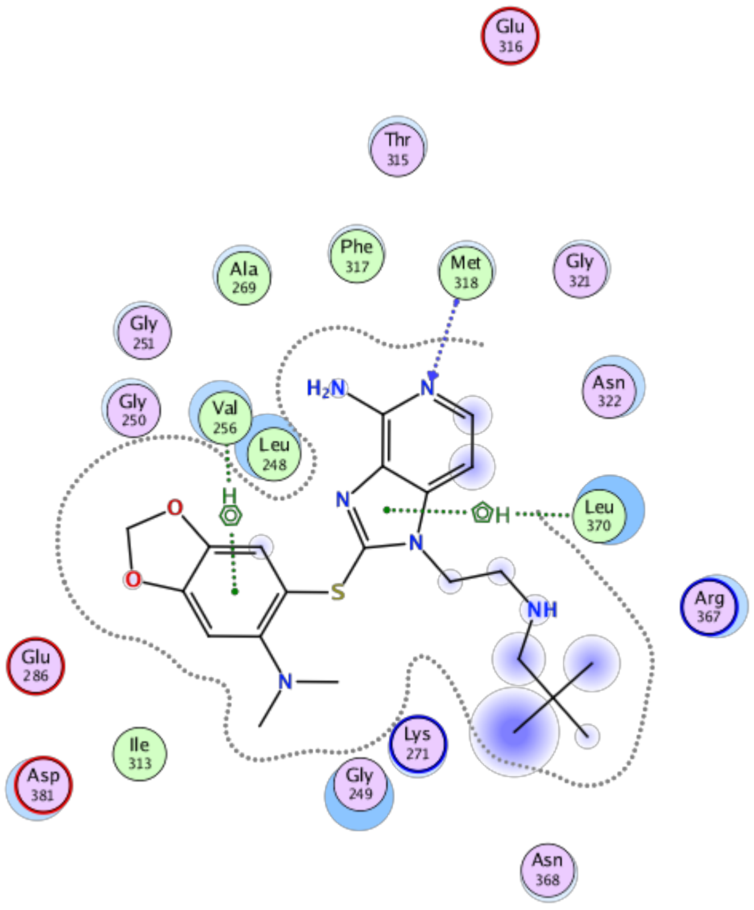

**Supplementary Figure 3.** Docking pose with the top MOE score (S = -8.73) for geldanamycin in ABL1 kinase. The MOE ligand interaction tool was used to generate the schematic diagram of protein-ligand interactions.

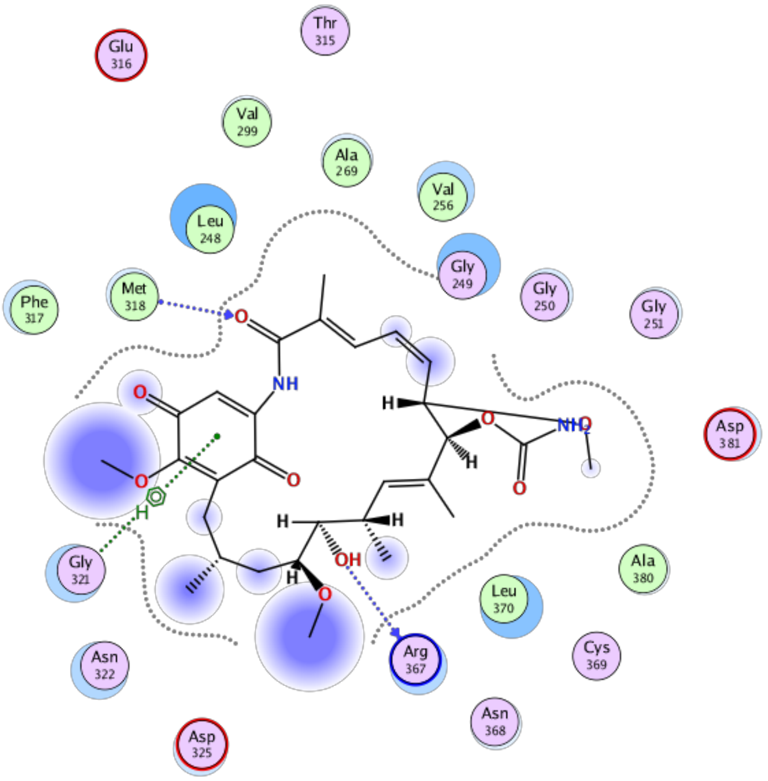

**Supplementary Figure 4.** Docking pose with the top MOE score (S = -6.94) for radicicol in ABL1 kinase. The MOE ligand interaction tool was used to generate the schematic diagram of protein-ligand interactions.

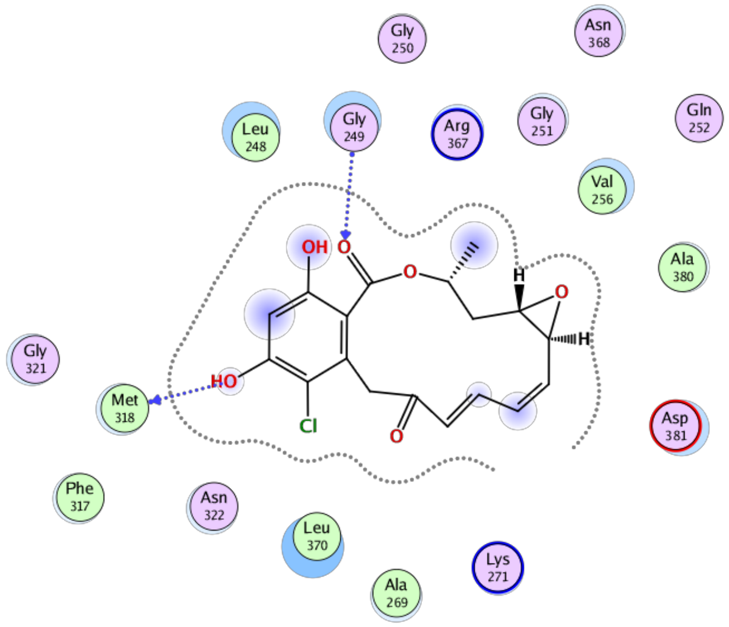
